# Prolonged Starvation Drives Epigenetic Remodeling: Insights from DNA Methylation Profiling in the Aquatic Pathogen *Flavobacterium columnare*

**DOI:** 10.1101/2025.10.01.679924

**Authors:** Yuxuan Zhang, Yanwen Shao, Shengnan Gao, Runsheng Li, Wenlong Cai

## Abstract

Nutrient scarcity is a common environmental stressor encountered by bacteria. Despite its significance in bacterial virulence and physiology, the underlying epigenetic mechanisms of bacterial adaptation to prolonged starvation remain poorly understood. In this study, we utilized the latest Nanopore R10.4.1 sequencing technology to comprehensively characterize the genome-wide methylation landscape of the freshwater fish pathogen *Flavobacterium columnare* following long-term starvation at different temperatures. Our results revealed significant methylation plasticity under starvation conditions, characterized by distinct motif-dependent patterns. Notably, demethylation of the 6mA-modified CAYNNNNNRTG motif emerged as a robust epigenetic signature of starvation adaptation, with functional enrichment of genes involved in pathways for translation and metabolism, suggesting the role of this methylase in regulating essential cellular functions under starvation stress. Additionally, another 6mA-modified GCAGA motif exhibited temperature-dependent variation, indicating its potential role in temperature-specific response. Together, this study provides novel insights into the epigenetic mechanisms underlying the bacterial adaptation to nutrient deprivation and establishes a valuable methodological reference for bacterial epigenetics analysis utilizing advanced Nanopore sequencing technologies.

## 1. Introduction

Bacteria constantly face environmental stressors, among which nutrient scarcity is particularly prevalent (Watson et al., 1998). Whether within host organisms or in their natural ecological niches, bacteria frequently endure extended periods of nutrient depletion, known as the “feast and famine cycle” (Holscher, 2021; Jørgensen and Boetius, 2007). This have driven the evolution of sophisticated survival strategies among bacteria, enabling them to manage nutrient shortages until environmental conditions improve (Zhu and Dai, 2023). For instance, some bacteria of *Bacillu**s*** spp. can form spores to endure prolonged nutrient deprivation, while non-spore-forming bacteria, such as *Bacillus subtilis* and *Micrococcus luteus* enter a dormant state or significantly reduce their growth rate (Gray et al., 2019; Kaprelyants and Kell, 1993). Prolonged starvation can also induce physiological changes, including alterations in cell and colony morphologies and enhanced antibiotic resistance (Finkel and Kolter, 1999; Harbi et al., 2013; Nguyen et al., 2011; Stretton et al., 1997). While previous studies have elucidated genetic variation and shrifts in gene expression patterns that contribute to bacterial adaption to starvation, the role of epigenetic regulation remains largely unexplored (Behringer et al., 2024; Hazan et al., 2021; Iyer et al., 2018; Switzer et al., 2018).

Some studies have linked DNA methylation with bacterial adaption to environmental stressors owing to its rapid and reversible regulatory characteristics (Riber and Hansen, 2021). As a prevalent epigenetic mechanism in prokaryotes, DNA methylation involves the transfer of methyl groups to specific DNA motifs by DNA methyltransferases (MTases), leading to the formation of modifications such as N6-methyl-adenine (6mA), C5-methyl-cytosine (5mC) and N4-methyl-cytosine (4mC) modifications. One of the most recognized functions of DNA methylation in bacteria is its integration into the restriction-modification (RM) system. This system serves as a defense against the invasion of foreign DNA, wherein MTases modify host DNA to prevent its degradation by restriction enzymes (Blow et al., 2016). Beyond its defensive role, DNA methylation is crucial for regulating a variety of physiological processes, such as gene expression and DNA mismatch repair (Casadesús and Low, 2006; Stone et al., 2023). While the role of DNA epigenetic modifications in stress responses has been extensively studied in eukaryotes, its significance in bacterial adaptation is increasingly being recognized (Houtepen et al., 2016; López et al., 2022). For example, methylation of the GATC motif has been associated with ciprofloxacin resistance in *Escherichia coli*, highlighting the role of DNA methylation in conferring antibiotic resistance (Cohen et al., 2016). Despite these insights, the broader landscape of methylation patterns and their underlying mechanisms that confer bacterial adaption under stress remains underexplored.

Advanced third-generation sequencing technologies, such as Single-Molecule Real-Time (SMRT) sequencing and Nanopore sequencing, have facilitated the *de novo* detection of bacterial methylation modifications. These methods directly identify methylation modifications during the sequencing process according to the differences in sequencing signals between nature DNA and unmethylated DNA (Beaulaurier et al., 2019). While SMRT sequencing has been widely applied in methylation detection, it faces challenges, such the requirement for high sequencing coverage and relatively low sensitivity in detecting 5mC modifications (Flusberg et al., 2010; Schatz, 2017). On the other hand, Nanopore sequencing is rapidly advancing in this field, supported by computational tools like Hammerhead (Liu et al., 2024a), Nanodisco (Tourancheau et al., 2021), and Tombo (Stoiber et al., 2017), which have been developed to analyze methylation from Nanopore data. However, these tools either lack the ability to distinguish between methylation types or depend heavily on motif contexts for modification identification (Amarasinghe et al., 2020; Tourancheau et al., 2021). In contrast, the optimized Dorado tool offers a significant improvement, enabling not only genome-wide identification of diverse methylation modifications but also quantification of their methylation levels. Additionally, it is worth noting that the recently emerging Nanopore R10.4.1 flowcell has achieved significant progress in improving the accuracy of methylation calling, providing a promising opportunity for high-accuracy DNA methylation detection (Ni et al., 2023). However, despite this advancement, Nanopore R10.4.1-based methylation analysis has rarely been applied to bacterial studies.

*Flavobacterium columnare* is the causative bacterium responsible for columnaris disease, which has led to substantial economic losses in global aquaculture due to its high mortality rate in various freshwater fish, such as catfish and tilapia (Amin et al., 1988; Burgos et al., 2023; Wagner et al., 2002). Columnaris is a warm water disease, generally occurring when water temperatures exceed 25 °C (Evenhuis et al., 2014). In aquaculture environments, *F. columnare* typically spreads through direct fish-to-fish contact and through waterborne transmission (Welker et al., 2005), and they could form biofilm as the pathogen reservoir (Cai et al., 2019; Declercq et al., 2019). Bacteria in these environments often experiences nutrient limitations, necessitating that *F. columnare* continuously adapt to starvation conditions to survive (Elser et al., 1995). Current research have shown that *F. columnare* can persist for extended periods in nutrient-depleted water without forming spores, accompanied by morphological changes and reduced virulence (Arias et al., 2012). Consistent with these findings, our preliminary research also demonstrated that *F. columnare* is capable of surviving in ddH₂O for over ten months. However, the molecular mechanisms driving its adaptation to prolonged starvation remain poorly understood.

In this study, we explored the epigenetic change of *F. columnare* in response to 10-month-long starvation. Since temperature affects the virulence and physiology of *F. columnare*, we also evaluated the impact of two environment-relevant temperatures (i.e., 22 and 28 °C) on the methylation profile after nutrient-starvation treatment. Nanopore R10.4.1 sequencing was applied to characterize DNA methylation in aquatic bacteria for the first time, enabling the identification and quantification of genome-wide DNA methylation at single-base resolution. Our study demonstrated that while the genome sequence of *F. columnare* remained highly conserved between nutrient-rich and nutrient-depleted conditions, its methylation patterns displayed motif-specific and temperature-tailored flexibility. Overall, this study provides novel insights into the methylation patterns underlying *F. columnare* adaptation to long-tern nutrient deprivation, offering potential avenues for improving infection control and mitigating the transmission of this pathogen in aquaculture systems. Importantly, our study provides a roadmap for the future practice of Nanopore sequencing in bacterial DNA methylation research.

## 2. Materials and Methods

### 2.1 Bacterial strain and starvation treatment

The *F. columnare* FCMF strain used in this experiment was isolated from mandarin fish (*Siniperca chuatsi*) infected with columnaris disease in Sichuan Province, China (Li et al., 2025). The bacteria were initially stored at -80°C in modified Shieh (MS) broth (i.g. 0.5% tryptone, 0.2% yeast extract, 45.6 µM CaCl_2_·2H_2_O, 1.08 mM KH_2_PO_4_, 1.2 mM MgSO_4_·7H_2_O, 4.1 µM FeSO_4_·7H_2_O, pH 7.2-7.4) containing 30% glycerol and were revived by growing for 24 hours in 5ml fresh MS broth at 28°C with agitation (125 rpm) (Cai and Arias, 2021). The culture was transferred into 35 ml of MS broth and incubated overnight under the same conditions (sample Fc-con). Aside from the transfer step, the culture medium was not changed throughout the process. The *F. columnare* cells were then centrifuged at 5000 rpm for 10 minutes, and the resulting pellets (∼3x10^9^ cells) were re-suspended in 10 ml of ddH₂O and maintained at 22°C and 28°C, respectively. The cells were maintained in ddH_2_O for ten months to maximize the contrast between nutrient-rich and nutrient-deficient conditions, with no replacement of the ddH₂O during this period. The two temperatures were selected because they reflect different growth dynamics for *F. columnare*: 28°C is the optimal growth temperature for *F. columnare*, while 22°C is the common room temperature. Due to the extremely poor DNA quality obtained from direct extraction of starved cultures, 1 ml of each ten-month-starved sample at 22 °C and 28 °C was transferred to 10 ml of MS broth for overnight culture (sample Fc-22 and Fc-28), prior to DNA extraction. This step was intended to improve DNA quality while minimizing the effect of DNA from dead cells.

### 2.2 DNA extraction, library construction and Nanopore sequencing

Genomic DNA of Fc-con, Fc-22, and Fc-28 was extracted using TaKaRa MiniBEST Universal Genomic DNA Extraction Kit (TaKaRa Bio Inc., Japan) after overnight culture. NanoDrop Spectrophotometer (Thermo Fisher Scientific, USA) and 1.5% agarose gel electrophoresis were used to assess the quality of the extracted DNA, prior to library preparation and sequencing. One µg of high-quality native DNA was used for whole-genome sequencing (WGS), following library preparation using the Native Barcoding Kit 24 V14 (SQK-NBD114.24, ONT, United Kingdom), according to the standard protocol. To establish the baseline methylation thresholds, whole-genome amplified DNA of Fc-con were employed as the unmethylated control. The whole genome amplification was performed using the REPLI-g Single Cell Kit (Qiagen, Germany), followed by library construction with the Native Barcoding Kit 24 V14 (SQK-NBD114.24, ONT). Sequencing of the prepared libraries was subsequently performed on the MinION platform (ONT), utilizing an R10.4.1 Flow Cell (FLO-MIN114, ONT).

### 2.3 Genome assembly and annotation

*De novo* genome assembly of Fc-con, Fc-22 and Fc-28 was performed by integrating the R10.4.1 simplex and duplex long reads. First, Dorado v0.8.3 (https://github.com/nanoporetech/dorado) was used to perform simplex and duplex basecalling on POD5 files generated from Nanopore whole-genome sequencing, with the <u>dna_r10.4.1_e8.2_400bps_sup@v5.0.0</u> model and <u>dna_r10.4.1_e8.2_5khz_stereo@v1.3</u> model, respectively. Samtools v1.9 (Danecek et al., 2021) was then used to extract the duplex reads. The quality of simplex and duplex reads was assessed using Giraffe v0.2.0 (Liu et al., 2024b). Prior to assembly, the ONT reads obtained by simplex basecalling were filtered using Filtlong with the parameters -min-mean-q 80 and -min_length 1000. The retained high-quality reads were assembled using Canu v2.2 (Koren et al., 2017) with the parameter genomeSize=3.4m. Additionally, the maxInputCoverage flag was set to 200 to randomly sample reads at a coverage depth of 200×. The assembled genome was subsequently polished using simplex reads with Medaka v1.11.3 (https://github.com/nanoporetech/medaka), followed by further polishing using duplex reads with Hammerhead v0.2.0 (Liu et al., 2024a). Finally, Dnaapler v0.7.0 (Bouras et al., 2024) was employed to reorient the chromosomes, ensuring they started from the DnaA gene. The completeness of polished assemblies was evaluated by BUSCO (Benchmarking Universal Single-Copy Orthologs) v5.7.1 (Manni et al., 2021), using 733 orthologs from the flavobacteriales_odb10 database as reference. To evaluate genome recovery rate and depth, the Nanopore long reads were aligned to the assembled genomes by minimap2 v2.28 (Li, 2018). All the sequencing data and genome assemblies are available on NCBI BioProject PRJNA1247036.

The genome assembly was functionally annotated using the Prokka v1.14.6 (Seemann, 2014), with Prodigal v2.6.3 (Hyatt et al., 2010) used for gene prediction. EggNOG-mapper v2.1.12 (Cantalapiedra et al., 2021) and InterProScan v5.69-101.0 (Jones et al., 2014) were used for Gene Ontology (GO) annotation. The promoter regions in the genome were predicted using PePPER v2.0 (de Jong et al., 2012), and the insertion sequence (IS) elements were predicted using ISEScan v1.7.2.3 (Xie and Tang, 2017).

### 2.4 Variant calling

To assess the role of genetic variation in *F. columnare* starvation tolerance, we provided a comprehensive view of SNPs and INDELs (insertions and deletions) in Fc-22 and Fc-28. To ensure a robust analysis, both assembly-based and read-mapping-based approaches were employed. In the assembly-based approach, the assemblies of Fc-22 and Fc-28 were aligned to the Fc control assembly using minimap2 v2.28, with Bcftools v1.20 (Li, 2011) and SyRI v1.7.0 (Goel et al., 2019) employed to detect SNPs and INDELs, respectively. For the read-mapping-based approach, sequencing reads from Fc-22 and Fc-28 were mapped to the Fc control assembly using Minimap2, followed by Bcftools-driven SNP and INDEL identification, with ploidy set to 1. To enhance the reliability of variant calls, only those with quality scores > 30 were retained. Additionally, the same read-mapping-based approach was applied to identify SNPs and INDELs between the Fc-con assembly and its reads. These regions were excluded from SNP and INDEL identification in Fc-22 and Fc-28 to minimize potential biases caused by inaccuracies in Fc-con assembly,.

### 2.5 High-confidence methylated site and motif identification

The Nanopore sequencing datasets were used to determine genome-wide methylation profiles for *F. columnare* under different nutritional stress conditions. Dorado was employed for modification basecalling of Nanopore reads, with the configuration of <u>dna_r10.4.1_e8.2_400bps_sup@v5.0.0_6mA@v2</u> and <u>dna_r10.4.1_e8.2_400bps_sup@v5.0.0_4mC_5mC@v2</u>. The resulting Nanopore reads with modification information were then mapped to the genome of Fc-con by Minimap2. To quantify methylation modifications, we leveraged the “pileup” command of Modkit v0.4.1 (https://github.com/nanoporetech/modkit) to aggregate 4mC, 5mC, and 6mA modification calls across all genomic positions using the “pileup” command. The 10% lowest confidence modification calls were excluded from counting to enhance accuracy. The methylation level of a specific methylation modification at each genomic site was defined as the “percent modified” output from the “pileup” command of Modkit software, calculated using the following formula:

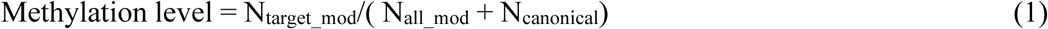

where N_target_mod_ was the number of calls classified as a residue with a target modification, N_all_mod_ was the number of calls classified as modified, and N_canonical_ was the number of calls classified as unmodified.

Given the noise in Nanopore sequencing, we utilized the whole genome amplification (WGA) dataset, which was practically free of methylation modifications, as a negative control to filter out spurious base modifications detected in Fc-con, Fc-22, and Fc-28 WGS dataset. Based on the methylation levels displayed at each site in both WGA and WGS sequencing, we attempted to determine high-confidence methylated sites using two different methodologies. In the first approach, we established thresholds of methylation levels for 6mA, 5mC, and 4mC modifications exclusively using the WGS dataset, simulating a scenario in which WGA data were unavailable. The threshold values were selected to maintain a false discovery rate (FDR) below 0.01. The false discovery rate was calculated using the following formula:

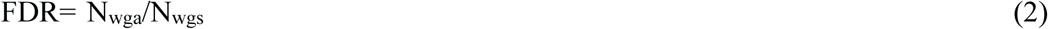

where FDR was the false discovery rate at this cutoff, N_wgs_ was the number of sites with methylation levels exceeding the cutoff in WGS sequencing, and N_wga_ was the number of sites with methylation levels exceeding the cutoff in both WGS and WGA sequencing. The second method involved defining thresholds based on the difference in methylation levels between WGS and WGA at each genomic site. This approach aimed to exclude genomic positions exhibiting similarly methylation signals in both datasets, thereby identifying genuine methylation events in the WGS samples.

The high-confidence methylated sites were used for subsequent methylated motif identification. Using BEDTools v2.31.1 (Quinlan and Hall, 2010), we extracted flanking regions around 6mA, 5mC, and 4mC sites (-12 bp and +12 bp), respectively. STREME from the MEME suite v5.5.7(Bailey et al., 2015) was then used to detect significantly enriched motifs in these sequences (*p*-value < 0.05). In parallel, methylation motif detection was also performed using the “motif search” function in Modkit software, with –min-sites set to 100. To further validate the identified motifs, we attempted to associate them with DNA methyltransferases (MTases). DNA methyltransferase prediction was conducted by performing BLASTP v2.15.0 (Camacho et al., 2009) against DNA methyltransferases in REBASE (Roberts et al., 2003) to identify the best hits with known recognition motifs, using the following criteria: identity >30%, e-value < 10⁻^10^, and bit score > 100. As a complementary approach, Nanomotif was also utilized to assign DNA methyltransferases to the identified motifs. To avoid potential false positives in the final methylation motif catalog, we systematically compared the methylation levels of each motif type between WGS and WGA datasets, ensuring a significant difference by the Wilcoxon Signed-Rank Test. Also, Hammerhead was used to calculate and compare the difference indices of each motif between WGS and WGA datasets.

### 2.6 Comparison of methylation levels under nutrient-deficient and nutrient-rich conditions

To understand the methylation adaptation of *F. columnare* to prolonged starvation, we analyzed its methylation levels under both nutrient-deficient and nutrient-rich conditions to investigate the trends of demethylation and methylation. Our analytical approach involved two levels of examination. Initially, we conducted a motif-level analysis, comparing methylation levels of each motif under the contrasting nutritional conditions. Recognizing the motif-driven nature of bacterial methylation, our analysis focused exclusively on high-confidence sites located within previously identified motifs to minimize the risk of false positives. Additionally, we restricted our examination to methylated sites consistently present across Fc-con, Fc-22, and Fc-28, to increase the robustness. We employed the Wilcoxon signed-rank test for this statistical analysis to determine significant differences. Subsequently, single-base-resolution differential methylation levels were also examined at these sites using the “dmr” command in Modkit, which calculated MAP-based *p*-values for each site. Sites exhibiting a MAP-based *p*-value < 0.05 and a methylation level change > 10% were considered significantly differentially methylated sites. This dual-level approach enabled a comprehensive understanding of the methylation changes in response to nutrient availability.

### 2.7 Functional characterization of methylation pattern changes under long-term nutrient deficiency

To further investigate the functional implications of methylation pattern changes, the significantly differentially methylated sites were mapped across the genome to determine their localization within CDSs or intergenic regions. GO enrichment analysis was performed on genes containing significantly differently methylated motifs using the clusterProfiler package v4.10.051 (Yu et al., 2012) in R.

## 3. Results

### 3.1 Morphological changes of *Flavobacterium columnare* colonies induced by long-term starvation

After 10 months of starvation, Fc-con, Fc-22 and Fc-28 was cultured on MS broth to evaluate the viability cells. The result showed that bacterial cells at both temperature were viable and revived on MS broth at a concentration of about 1x10^3^ CFU/ml. Interestingly, these bacteria under this extended period of nutrient starvation displayed striking differences in colony morphology compared to Fc-con. The colonies of Fc-con under nutrient-rich conditions exhibited a flat, root-like structure. In contrast, the nutrient-deprived Fc-22 and Fc-28 displayed identical colony morphologies, featuring rounded edges and smooth surfaces with significantly decreased cellular size (Supplementary Fig. S1).

### 3.2 Genome sequencing, assembly, and annotation of *F. columnare*

This study sequenced *F. columnare* under different nutritional conditions by utilizing Nanopore sequencing technology. High-quality genomic DNA was extracted from Fc-con, Fc-22, and Fc-28, with concentrations exceeding 40 ng/μL, A260/A280 ratios above 1.8, A260/A230 ratios above 2.0, and apparent integrity on 1.5% agarose gels (Supplementary Table S1 and Supplementary Fig. S2). The Nanopore sequencing generated 665.6 Mb, 788.4 Mb, and 1.3 Gb of data for Fc-con, Fc-22, and Fc-28, respectively. The average read lengths were 4,773 bp, 3,428 bp, and 989 bp, with N50 values of 14,777 bp, 9,747 bp, and 1,603 bp (Supplementary Table S1). These differences likely resulted from technical or systematic biases introduced post DNA extraction. Their read accuracies exceeded 97% estimated by Giraffe. The total duplex read output for Fc-con, Fc-22, and Fc-28 were 19.3 Mb, 16.5 Mb, and 14.5 Mb, respectively. Additionally, whole-genome amplification sequencing of Fc-con produced total bases of 1.9 Gb, with an average read length of 888 bp utilizing Nanopore sequencing technology.

*De novo* assembly of Fc-con, Fc-22, and Fc-28 resulted in circular genomes with highly similar sizes, all approximately 3.36 Mb (Fig. 1a and Table 1). A subsequent round of polishing with duplex reads corrected 23, 33, and 18 bases in Fc-con, Fc-22, and Fc-28, respectively. In the final assemblies of Fc-con, Fc-22, and Fc-28, we identified 725 (99.7%), 724 (99.6%), and 724 (99.6%) complete Benchmarking Universal Single-Copy Orthologs (BUSCOs) from flavobacteriales_odb10, demonstrating a high level of completeness. Furthermore, by mapping Nanopore reads back to the genomes, we confirmed genome recovery rates of 100%, indicating high assembly accuracy. The genome annotation identified 2894, 2892, and 2898 coding sequences (CDSs) in Fc-con, Fc-22, and Fc-28 assemblies.

**Figure 1.**
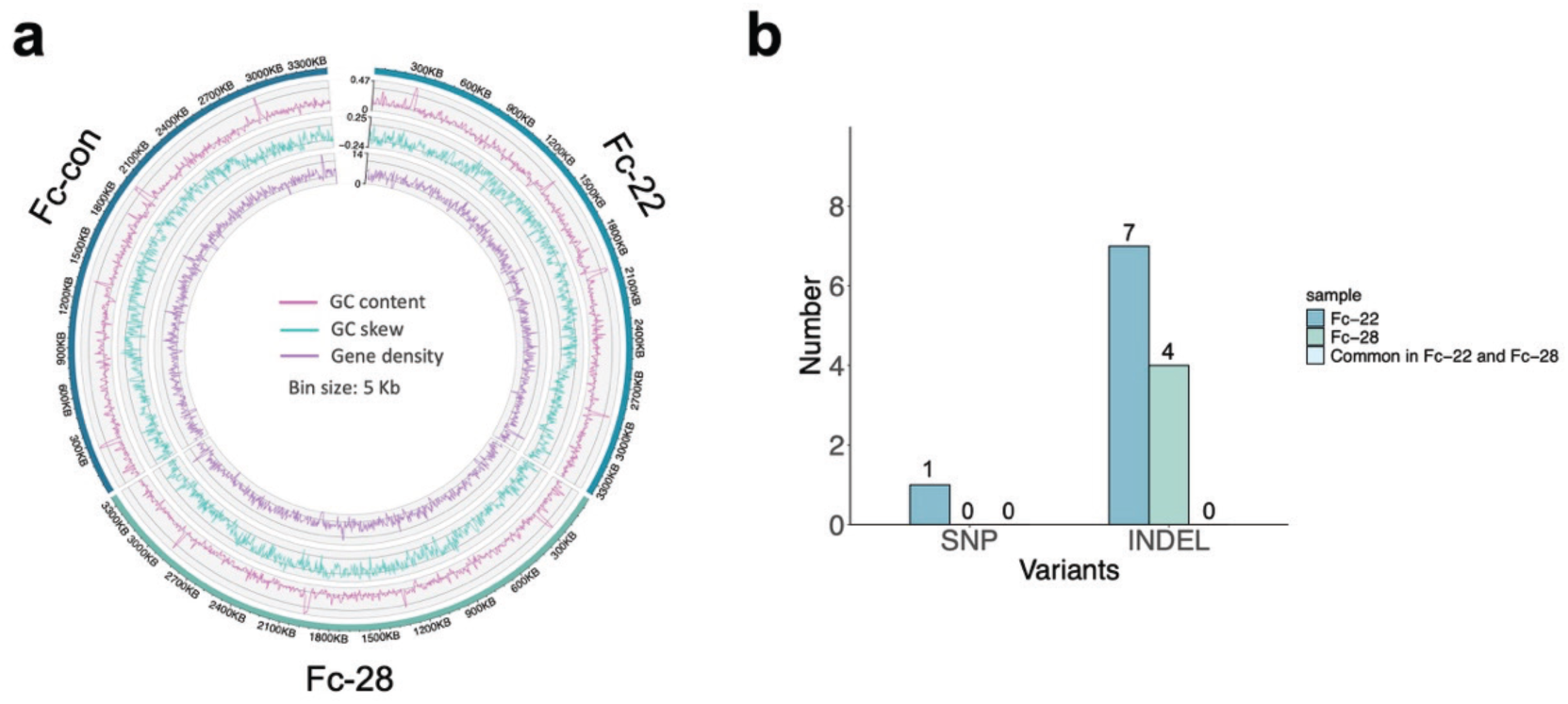
Conserved genomic sequence of *F. columnare*. (**a**) Circos plot of Fc-con, Fc-22, and Fc-28. For inner to outer layers: gene density, GC skew, and GC content. (**b**) Number of SNPs and INDELs in Fc-22, Fc-28, and shared in both samples. The variations are identified using the read-mapping-based method.

**Table 1.** Statistics of *F. columnare* assembly and annotation.

|  | Fc-con | Fc-22 | Fc-28 |
| --- | --- | --- | --- |
| Assembly size (bp) | 3,365,019 | 3,365,936 | 3,358,135 |
| GC content (%) | 31.63 | 31.63 | 31.63 |
| Complete BUSCOs | 725 (99.7%) | 724 (99.6%) | 724 (99.6%) |
| Missing BUSCOs | 1 (0.1%) | 1 (0.1%) | 1 (0.1%) |
| Genome recovery rate (%) | 100 | 100 | 100 |
| Predicted CDSs | 2,894 | 2,898 | 2,892 |
| Mean CDS length (bp) | 996 | 995 | 995 |
| Promoters | 983 | 983 | 983 |
| IS | 68 | 67 | 67 |

### 3.3 Few genetic variations occurred in *F. columnare* subjected to long-term starvation

To investigate the potential genetic backgrounds associated with long-term starvation in *F. columnare*, we identified genetic variants in the two starved bacteria, Fc-22 and Fc-28 by comparing to Fc-con. The read-mapping approach detected only one non-synonymous SNP (Val127Phe) in Fc-22, while 7 and 4 unique INDELs were identified in Fc-22 and Fc-28, respectively, with 2 and 1 INDELs located in intergenic regions (Fig. 1b and Supplementary Table S2). Parallel analysis using the assembly-based approach identified 26 SNPs in Fc-22 and 24 SNPs in Fc-28, of which 20 and 14 SNPs were located in the CDS regions, with 9 and 7 being synonymous (Supplementary Table S2). Only 3 overlapping SNPs were found, none of which were within CDS regions. Additionally, 13 and 12 INDELs were identified in Fc-22 and Fc-28, including only 2 overlapping INDELs. Overall, based on the integrated results from both methods, few genomic regions in *F. columnare* exhibited variations under nutrient-deficient conditions at both temperature, with almost no INDELs and SNPs conserved across both temperatures, implying that such a limited number of variants could be possibly attributed to random sequencing errors. Moreover, many identified variants were predicted to have minimal functional impact on protein sequences. Consequently, we speculated that starvation adaption of *F. columnare* was likely not the result of genetic variation.

### 3.4 High-confidence methylated site identification

Research has increasingly emphasized the role of DNA epigenetic regulation in conferring bacterial stress adaption. To investigate the association between starvation endurance and DNA epigenetic modifications in *F. columnare*, we first utilized Nanopore sequencing technology to uncover its genome-wide methylation landscape. Using the recently developed Dorado modification-calling models, we analyzed the native DNA from Fc-con, Fc-22, and Fc-28, detecting extensive methylation signals of 6mA, 5mC, and 4mC across over 2.5 million genomic sites (Supplementary Table S3).

To evaluate possible false positives in the detection of DNA modification, we compared the Dorado prediction result from WGA sequencing data using amplified unmodified DNA, to the WGS sequencing data using the native bacterial DNA. A substantial proportion of these detected modification sites exhibited comparable methylation levels between WGS and WGA result, suggesting a possibly high level of false positives when replying solely on WGS ONT sequencing for methylation profiling (Fig. 2a and Supplementary Fig. S3a,b). Moreover, the overall methylation levels between WGA and WGS sequencing were also close, especially for 4mC modifications (Supplementary Fig. S4). The background noise in Dorado WGS results needed to be filtered before further DNA methylation analysis.

**Figure 2.**
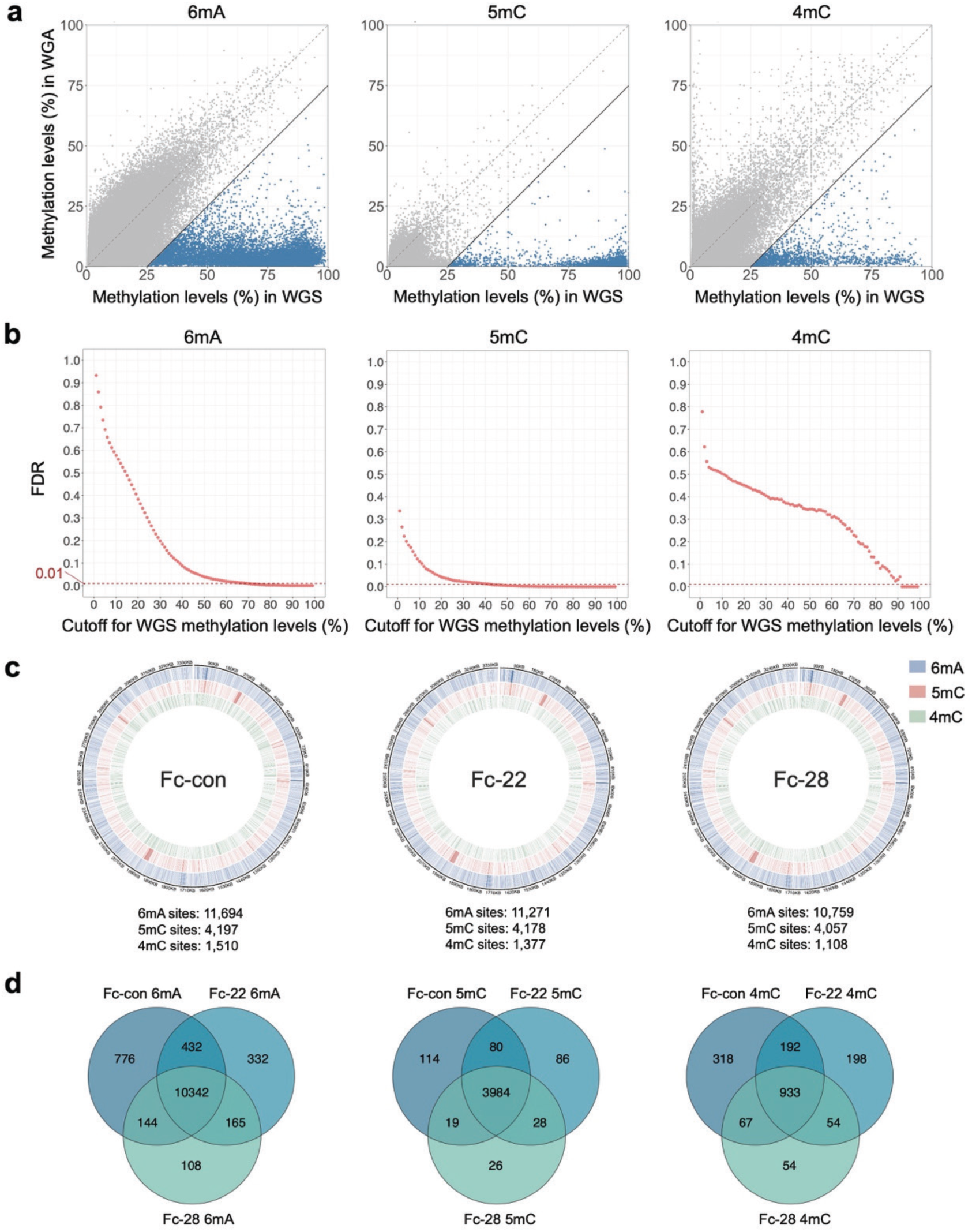
Filtering of raw base modifications in the native DNA of Fc-con output by Dorado. (**a**) Filtering based on the difference in methylation levels between WGS and WGA sequencing. The points represent individual base modifications. Most base modifications are distributed around the diagonal, indicating highly similar methylation levels between WGS and WGA. The blue points represent high-confidence base modifications filtered using the criterion "WGS methylation level - WGA methylation level > 25%". (**b**) False discovery rates (FDR) under filtering based solely on methylation levels in WGS sequencing. Methylation levels required for FDR < 0.01 are significantly high. (**c**) Circos plots for high-confidence 6mA, 5mC, and 4mC methylation sites in Fc-con, Fc-22, and Fc-28, filtered using the criterion "WGS methylation level - WGA methylation level > 25%". (**d**) Venn diagrams showing the overlap of high-confidence 6mA, 5mC, and 4mC methylation sites in Fc-con, Fc-22, and Fc-28.

To effectively eliminate these false positives, we sought to define stringent methylation thresholds based on comparative analysis between WGS and WGA datasets. Initially, we identified methylation level thresholds associated with false discovery rates below 0.01. For the 6mA modifications, methylation levels above 70% were required to ensure high confidence, and for 5mC modifications, this threshold was approximately 43% (Fig. 2b, Fig. S2c,d, and Supplementary Table S4-5). However, applying such strict criteria risked excluding biologically significant methylated sites with moderate methylation levels. For 4mC modifications, a methylation level cutoff of at least 94% was required to achieve acceptable false discovery rates, but this stringent threshold drastically reduced the number of retained 4mC sites to fewer than five (Fig. 2b, Fig. S2c,d, and Supplementary Table S6). Considering the limitations of setting thresholds based solely on methylation levels from WGS sequencing, we refined our filtering strategy by integrating differential methylation analysis between WGS and WGA sequencing data. Specifically, we defined a high-confidence methylation site as one where the methylation level in WGS exceeded that in WGA by at least 25% (i.e., WGS methylation level – WGA methylation level > 25%), which effectively eliminated the vast majority of background noise (Fig. 2a, Supplementary Fig. S3a,b).

Following the refined cutoff, we identified high-confidence methylation sites, including 11,694 6mA-modified, 4,197 5mC-modified, and 1,510 4mC-modified sites in Fc-con; 11,271 6mA-modified, 4,178 5mC-modified, and 1,377 4mC-modified sites in Fc-22; and 10,759 6mA-modified, 4,057 5mC-modified, and 1,108 4mC-modified sites in Fc-28, respectively (Fig. 2c). Remarkably, among these sites, approximately 90% were consistently observed across all conditions in *F. columnare* (Fig. 2d). The high consistency suggest the reliability of our identification of methylated sites.

### 3.5 Methylated motifs were conserved under different conditions

Methylated motif enrichment was performed on the detected high-confidence methylated sites. Utilizing both STREME and “motif search” function of Modkit, we identified 8 distinct methylated motifs: C6mAYNNNNNRTG, GAA6mANNNNNNNNTGG, GA6mATTC, GC6mAGA, ATCG6mAT, CAT6mATG, C5mCNGG, and CGTA4mCG (Fig. 3a, and Supplementary Table S7-9). Among these motifs, six were associated with the 6mA modification, while the other two corresponded to 5mC and 4mC modifications. The analysis revealed that the GCAGA motif was the most prevalent, identified at over 4000 methylated sites across the genome, while the CATATG motif was the least abundant (Fig. 3b and Supplementary Table S9). The majority of methylated motifs were localized within CDS regions, with CDS usage rates around 90% (Supplementary Fig. S5a,b). Functional analysis indicated a notable localization of these methylated motifs within CDS of genes categorized under the COG categories of Replication, recombination and repair (L), Cell wall/membrane/envelope biogenesis (M), and Amino acid transport and metabolism (E) (Supplementary Fig. S5c). Gene Ontology (GO) enrichment analysis corroborated these findings, demonstrating significant representation of methylated motifs within genes implicated in critical cellular processes such as DNA replication and tRNA aminoacylation pathways (Supplementary Fig. S5d). This suggests that these methylated motifs may play fundamental roles in the biological processes of *F. columnare*.

**Figure 3.**
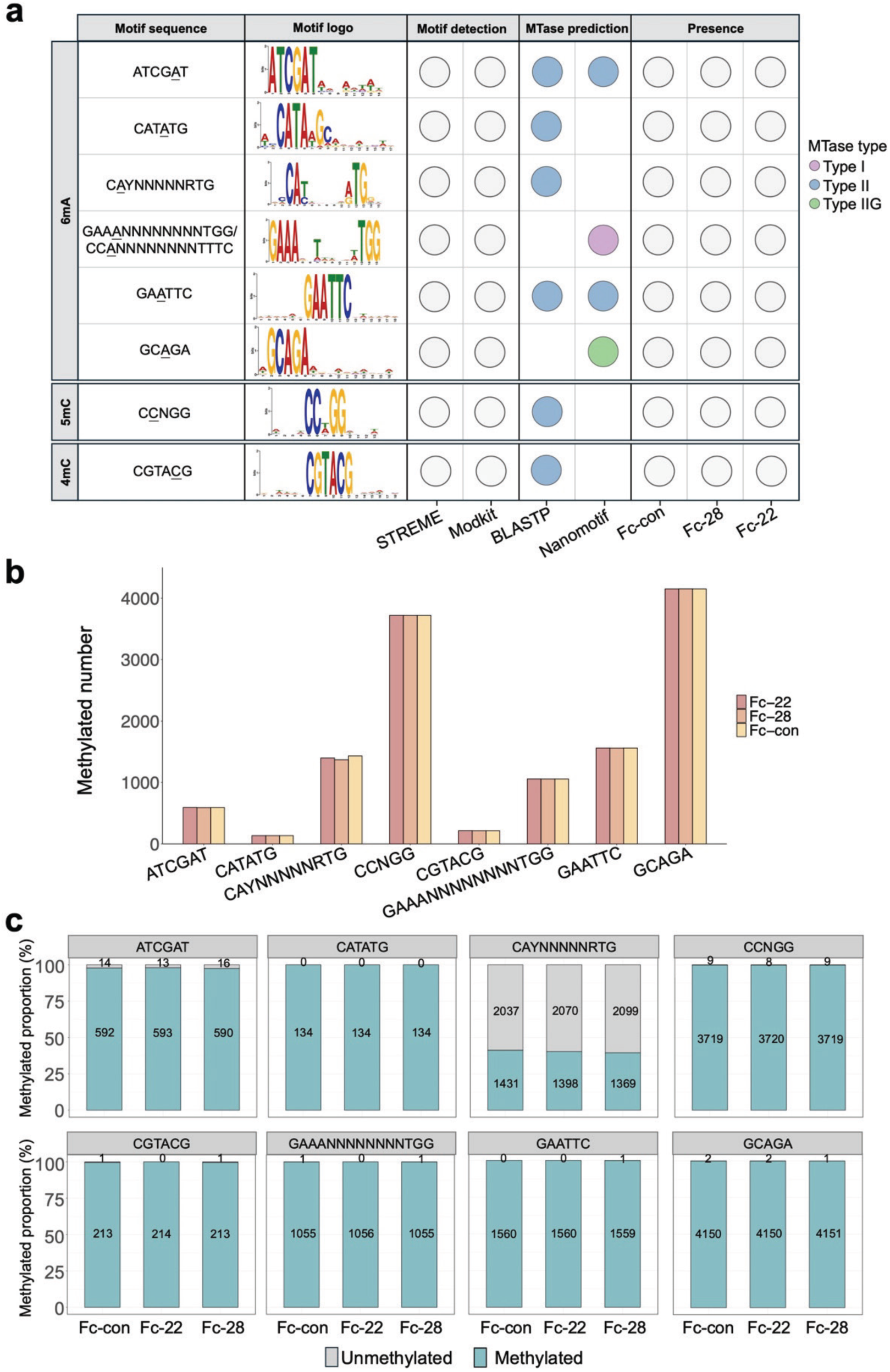
Conservation of methylated motifs in *F. columnare* under nutrient-rich and nutrient-deprived conditions. (**a**) Methylated motifs identified in Fc-con, Fc-28, and Fc-22 based on high-confidence methylated sites. The underscores beneath the motif sequences indicate the positions of methylation. The motif sequences are determined by integrating results from STREME, Modkit motif enrichment, and DNA methyltransferase identification. The motif logos are represented only by STREME enrichment results of Fc-con, and only the logo from one strand is displayed for each motif. The circles indicate that methylated motif were successfully identified or assigned to a methyltransferase using different methods, or that they were present in Fc-con, Fc-28, or Fc-22. (**b**) Methylated numbers of each motif in Fc-con, Fc-22, and Fc-28. Only the sequence from one strand is displayed for non-palindromic bipartite motifs. (**c**) Methylated proportion across the whole genome of each motif. The numbers on the bars show the methylated number and unmethylated number.

To confirm the authenticity of these methylated motifs, we compared their methylation levels and difference indices in native WGS datasets to those obtained from WGA control (Supplementary Fig. S6). As expected, methylation levels and differential indices were significantly higher in WGS datasets compared to WGA controls, supporting the authenticity of our identified motifs. Furthermore, we successfully assigned putative DNA methyltransferases to each identified motif through BLASTP searches against REBASE and through the Nanomotif tool, providing additional validation and strengthening confidence in our motif-identification pipeline (Supplementary Table S10). Overall, there were one Type I methyltransferase, one Type IIG methyltransferase, and six Type II methyltransferases, across the conditions tested. Among the high-confidence methylation sites identified in Fc-con, Fc-22, and Fc-28, around 80% were located within these identified motifs (Supplementary Fig. S7), confirming the comprehensiveness of our motif identification and the rationality of our filtering criteria for raw detected methylated sites. However, the motif coverage for 4mC-modified sites was markedly lower, constituting less than 20% of 4mC sites in all conditions (Supplementary Fig. S7). According to STREME enrichment results, many of these high-confidence 4mC-modified sites overlapped motifs typically associated with 6mA modifications, suggesting the possibility of confounding signals due to adjacent 6mA methylation events (Supplementary Table S7).

It is noteworthy that all of the motifs were consistently detected across different conditions, demonstrating highly conserved genome-wide methylation occurrence. Their methylated sites showed nearly consistent distribution across major genomic features, including promoters, CDS, and insertion sequences (IS), under different conditions (Supplementary Table S11). Additionally, most of the motifs showed near-universal methylation across the genome in all conditions, with methylation occurrence rates reaching nearly 100% (Fig. 3c and Supplementary Table S9). The exception was observed only for CAYNNNNNRTG, which showed a methylation occurrence of only around 40%. However, despite this lower methylation occurrence, the spatial consistency of its methylation across the genome remained remarkably high across the tested conditions (Supplementary Table S9).

### 3.6 Methylation levels were downregulated under prolonged starvation, particularly in CAYNNNNNRTG motif

In the case of high conservation of methylated motifs, we further analyzed their methylation changes (i.e. demethylation and methylation) to uncover specific patterns of methylation changes occurring in *F. columnare* under starvation stress. Focusing on the conserved methylated sites within these motifs, we discovered the plasticity in methylation levels of *F. columnare* under starvation conditions. Overall, these sites exhibited high methylation levels in different conditions, with the majority exceeding 50% (Fig. 4a). However, nutrient-deficient conditions led to an overall downregulation in methylation levels, indicating a trend toward demethylation (Fig. 4a).

**Figure 4.**
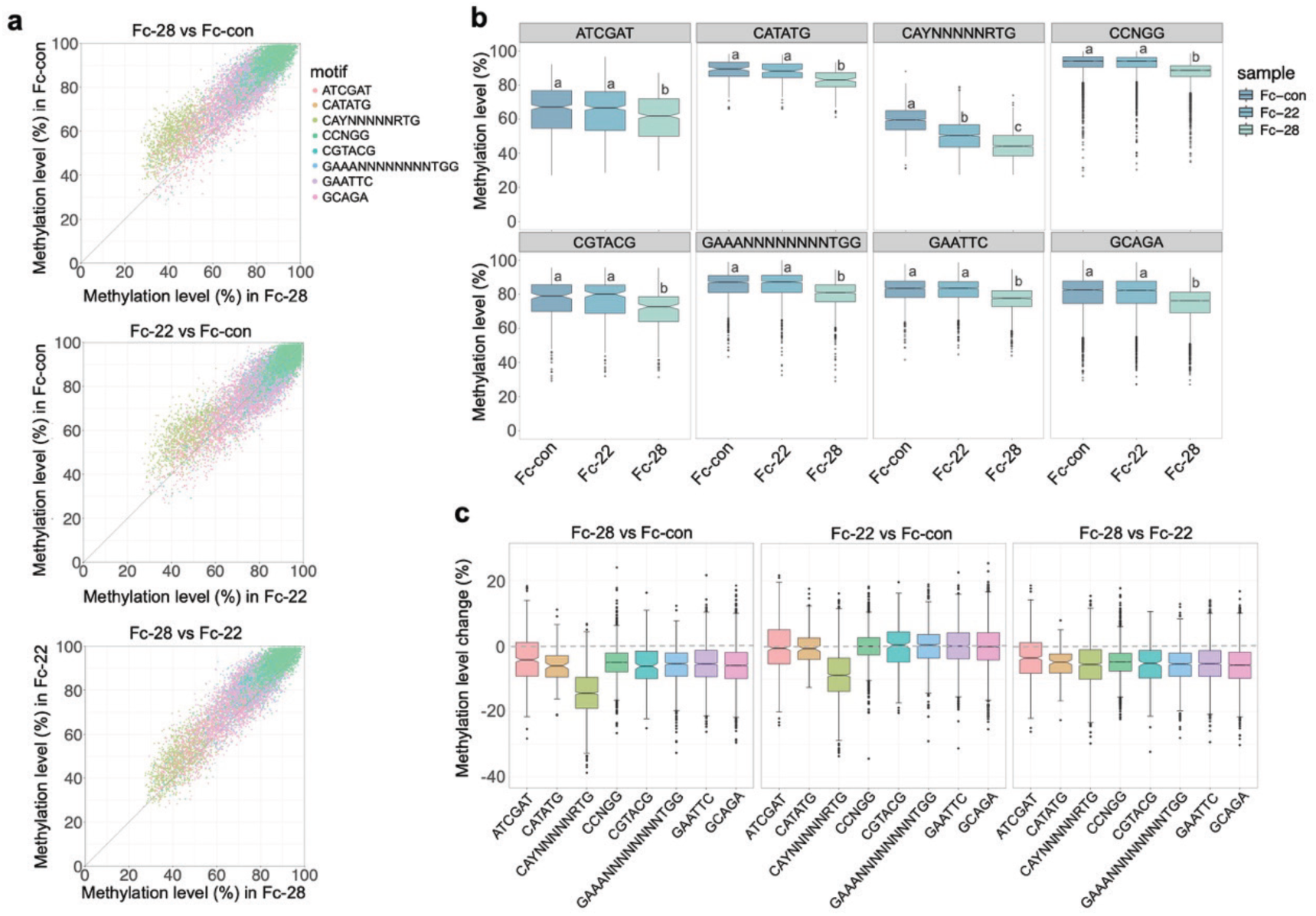
Downregulation of methylation levels at motif-level resolution in starved *F. columnare*, especially for the CAYNNNNNRTG motif. (**a**) Dot plots showing the methylation levels of each motif-associated methylated site under different conditions. (**b**) Box plots comparing the methylation levels of methylated sites for each motif between different conditions. Different letters above the boxes indicate significant differences (Wilcoxon signed-rank test *p* < 0.01). (**c**) The magnitude of changes in methylation levels of motif-associated methylated sites for Fc-28 compared to Fc-con, Fc-22 compared to Fc-con, and Fc-28 compared to Fc-22.

Notably, despite the consistent demethylation trend, methylation responses differed between the two starvation temperatures assessed. At 28°C, nutrient deprivation resulted in a significant overall decrease in methylation levels across all identified motifs (*p* < 0.01; Fig. 4b). In contrast, at 22°C, methylation levels remained relatively stable for most motifs, with a significant reduction observed exclusively in the CAYNNNNNRTG motif (*p* < 0.01; Fig. 4b). Interestingly, the CAYNNNNNRTG motif also exhibited the greatest magnitude of methylation level reduction under starvation conditions at both temperatures, which suggested its sensitivity to nutrient stress (Fig. 4c). However, when comparing methylation level changes between the two starvation temperatures (22°C and 28°C), the magnitude of change within the CAYNNNNNRTG motif was comparable to that observed in other motifs (Fig. 4c). Importantly, these results are highly consistent with those from Hammerhead (Supplementary Figure S6b). Specifically, the CAYNNNNNRTG motif consistently showed a significant decrease in difference index under starvation conditions, while the other motifs exhibited significant reductions only under starvation at 28 °C except for the CATATG motif, and remained stable under starvation at 22 °C. These observations suggest the potential importance of CAYNNNNNRTG motif in adaptation to starvation stress. Functional annotation of the 1,350 methylation sites associated with the CAYNNNNNRTG motif revealed that 1,252 sites were localized within genes significantly implicated in replication, tRNA aminoacylation, and siderophore-mediated iron transmembrane transport (Supplementary Fig. S5c-d).

### 3.7 CAYNNNNNRTG-related methylation reduction within translation and metabolism genes may represent a key epigenetic mechanism for starvation adaption

Having observed an overall declining trend in methylation levels due to prolonged starvation, we performed single-base resolution differential analysis to reveal significantly altered sites. Our results further reinforced the critical role of CAYNNNNNRTG-associated demethylation events in the response to starvation stress.

We first profiled single-base-level methylation changes under each starvation condition, revealing both shared and unique epigenetic characteristics compared to nutrient-rich control. In Fc-28, 1,546 sites showed significantly reduced methylation levels, while only 7 sites exhibited significantly increased methylation levels (Fig. 5a). Similarly, in Fc-22, 220 sites were significantly downregulated, with 64 sites showing significant upregulation(Fig. 5b). Motif analysis revealed distinct patterns between the two starvation temperatures. In Fc-28, the significantly downregulated sites mainly corresponded to three motifs, CAYNNNNNRTG, CCNGG, and GCAGA, with each accounting for comparable proportions (Supplementary Fig. S8a). Conversely, in Fc-22, the majority of significantly downregulated sites were located with the CAYNNNNNRTG motif alone (Supplementary Fig. S8c). Since these significantly downregulated sites in Fc-28 and Fc-22 were overrepresented among 882 and 175 CDSs, we conducted further functional analysis to elucidate their biological implications (Supplementary Fig. S8b,d,e-f). GO analysis revealed that, under starvation at 28 ℃, genes with significant downregulated sites primarily participated in processes related to replication and translation, as evidenced by significantly enriched GO terms including “DNA replication”, “DNA primase activity”, “tRNA aminoacylation”, “tRNA aminoacylation for protein translation” and “amino acid activation” (Supplementary Fig. S8e). In contrast, at the lower starvation temperature of 22 °C, significant methylation level reductions was not only observed in translation-related genes but also genes involved in various metabolic processes, such as “cellular nitrogen compound metabolism”, “macromolecule metabolism”, and “amide metabolic processes” (Supplementary Fig. S8f). Additionally, genes containing significantly downregulated methylation sites within the CCNGG motif were significantly enriched in functions related to ribosome biogenesis and methyltransferase activity. Notably, separate GO enrichment analyses of significantly downregulated sites in Fc-28 and Fc-22 each independently revealed enrichment of genes related to the siderophore transmembrane transport process (Supplementary Fig. S8e,f).

**Figure 5.**
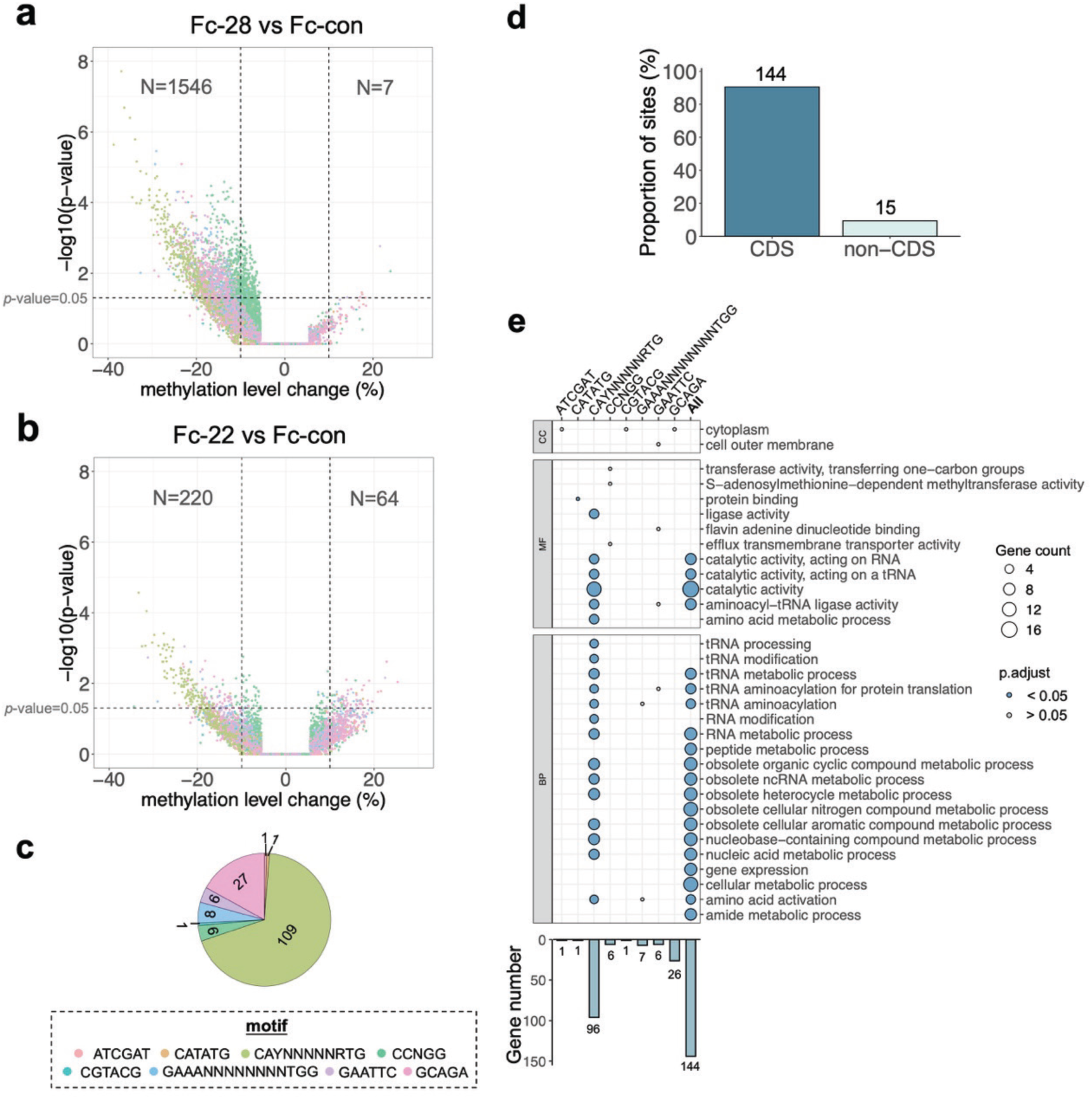
Significant downregulation of methylation levels at single-base resolution, predominantly featuring the CAYNNNNNRTG motif, in starved *F. columnare*. (**a-b**) Volcano plots showing sites with significant changes in methylation levels identified by single-base resolution differential methylation analysis in Fc-28 (a) and Fc-22 (b), compared to Fc-con. A *p*-value < 0.05 and a methylation level change of < -10% or > 10% were considered as significant downregulation or significant upregulation, respectively. Almost all the sites with significant changes are those downregulated. (**c**) Pie chart showing the number of significantly downregulated sites shared by Fc-28 and Fc-22, based on their distribution across different motifs. Most of the sites were associated with CAYNNNNNRTG. (**d**) CDS usage associated with significantly downregulated sites shared by Fc-28 and Fc-22. The numbers on the bars show the numbers of sites located within or not within CDSs. (**e**) Bar plot displays the number of CDSs containing the shared significantly downregulated sites for each motif. Dot plot shows the GO enrichment analysis of genes containing shared significantly downregulated sites for each motif. When more than 3 significant GO terms (Benjamini-Hochberg (BH)-adjusted *p*-value < 0.05) were identified, the top 20 are shown, and when fewer than 3 significant GO terms were identified, the top 3 are displayed. Blue and gray dots represent significant and non-significant enrichment, respectively.

To clearly identify conserved starvation-induced epigenetic responses across temperatures, we then focused specifically on overlapping significantly downregulated sites between Fc-28 and Fc-22 in relative to Fc-con. We identified 159 shared sites, the majority of which occurred within the CAYNNNNNRTG motif (Fig. 5c). GO enrichment analysis indicated that these conserved demethylated sites were predominantly associated with genes involved in translation, such as those encoding alanine-tRNA ligase, cysteine-tRNA ligase, and valyl-tRNA synthetase (Fig. 5d,e). Also, some of them were located in genes related to metabolic pathways, including those encoding S9 family peptidase and adenylosuccinate synthetase purA. Notably, nearly all of these enriched genes were attributed specifically to significantly downregulated sites within the CAYNNNNNRTG motif, highlighting the motif-specific functional implications under starvation stress. Collectively, these findings demonstrated that motif-dependent demethylation, particularly involving CAYNNNNNRTG, constituted a conserved and robust epigenetic signature of *F. columnare* in response to prolonged starvation. This targeted methylation remodeling within translation and metabolism genes likely played a critical role in survival under nutrient-limited conditions.

### 3.8 GCAGA-associated methylation exhibited temperature-dependent patterns on replication-related genes under starvation

Separate from the conserved starvation-associated epigenetic response described above, we further uncovered temperature-dependent methylation changes under the two starvation conditions, primarily involving a distinct motif, GCAGA. Specifically, by detecting differentially methylated sites between Fc-28 and Fc-22, we revealed that *F. columnare* exhibited predominantly lower methylation levels under starvation at 28 °C compared to 22 °C (Fig. 6a), which indicated a significant interplay between temperature and epigenetic adjustments. A total of 1,079 significantly downregulated sites were identified in Fc-28, most of which were associated with the GCAGA motif, while only 3 sites showed significantly increased methylation (Fig. 6b). These downregulated sites identified at 28 °C were predominantly located within 686 genes, significantly enriched in the DNA replication pathway, such as DNA primase *dnaG* and DNA polymerase III subunit beta gene (Fig. 6c,d). Importantly, significant demethylation specifically within the GCAGA motif contributed substantially to this enrichment. These observations indicated that the differential methylation, occurring particularly within the GCAGA motif, may represent a temperature-responsive epigenetic pattern related to DNA replication genes during adaption to starvation stress. Altogether, this suggested a motif-dependent, temperature-sensitive epigenetic mechanism that operated under starvation, distinct from the core starvation response characterized earlier.

**Figure 6.**
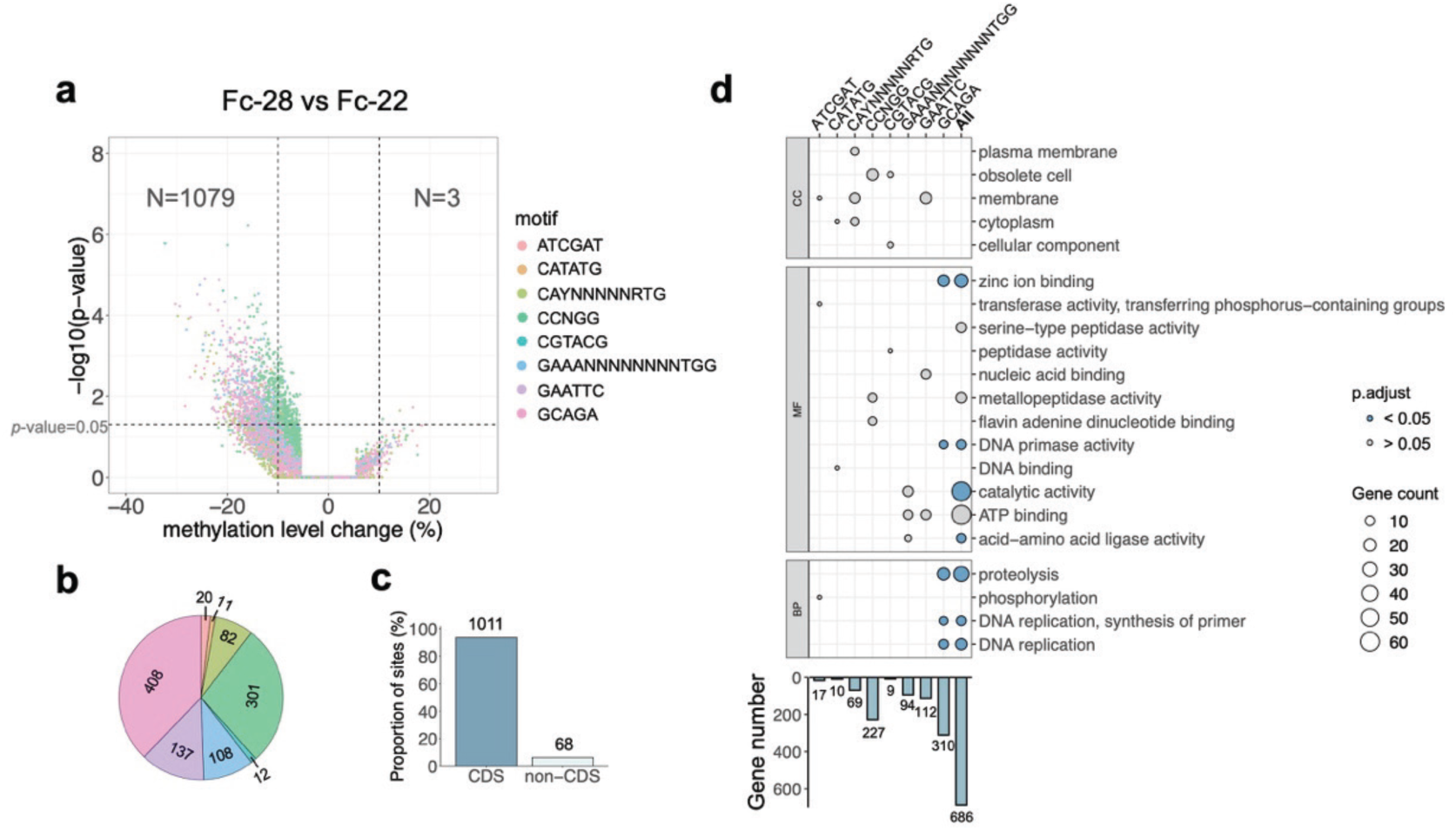
Significant differential methylation levels at single-base resolution, predominantly featuring the GCAGA motif, between starvation conditions at 28 °C and 22 °C. (**a**) Volcano plot showing sites with significant changes in methylation levels identified by single-base resolution differential methylation analysis in Fc-28, relative to Fc-22. (**b-c**) Proportion of motifs (b) and CDS usage (c) associated with significantly downregulated sites in Fc-28 compared to Fc-22. The numbers in the pie chart and on the bars indicate the corresponding site counts. (**d**) Bar plot displays the number of CDSs containing significantly downregulated sites for each motif in Fc-28. Dot plot illustrates the GO enrichment analysis of genes containing significantly downregulated sites for each motif in Fc-28. When more than 3 significant GO terms were identified, the top 10 are shown, and when fewer than 3 significant GO terms were identified, the top 3 are displayed. Blue and gray dots represent significant and non-significant enrichment, respectively.

## 4. Discussion

A longstanding question in microbiology is how bacteria adapt to frequently encountered environmental stresses to ensure their survival. Although multiple genetic and transcriptional factors involved in stress adaptation have been well characterized, there is increasing recognition that epigenetic mechanisms may also play a pivotal role (Riber and Hansen, 2021). Despite the fundamental importance of epigenetic regulation in modulating various bacterial processes, few studies have comprehensively characterized bacterial stress adaptation from an epigenetic perspective. Recent advances in third-generation sequencing technologies have now enabled comprehensive genome-wide characterization of bacterial methylation profiles. Among these, Nanopore R10.4.1 sequencing represents one of the most advanced DNA detection techniques available. In this study, we applied Nanopore R10.4.1 sequencing for the first time to an aquatic bacterium, thoroughly characterizing the genome-wide methylation patterns associated with the adaptation of *F. columnare* under prolonged starvation stress.

### 4.1 Lack of genetic variation suggests epigenetic mechanisms in *F. columnare* starvation persistence

Bacterial persistence under stress is generally considered as a fully reversible phenomenon, commonly thought to be regulated by changes in regulation of gene expression rather than by genetic mutations (Riber and Hansen, 2021). In this study, we found that the freshwater fish pathogen *F. columnare* can persist for up to ten months in nutrient-depleted conditions. To investigate the underlying adaptive mechanisms, we first assessed genetic variation in *F. columnare* following prolonged starvation treatment. Both assembly-based and read-mapping-based analysis revealed an extremely limited low number of SNPs and INDELs, with the possibility that these variants could be attributed to sequencing errors. Consequently, our results strongly suggest that the bacterium persistence observed in *F. columnare* was a consequence of epigenetically regulated mechanisms.

### 4.2 Reliable identification of Genome-wide methylated sites and motifs

Nanopore R10.4.1 sequencing, coupled with the downstream Dorado analysis, exhibits remarkable superiority in achieving whole-genome methylation profiling with single-base resolution, which has scarcely been implemented in bacterial contexts. Nevertheless, we observed considerable noise in the raw output from Dorado, which complicated accurate base modification identification in the absence of whole-genome amplification (WGA) sequencing data, especially for 4mC modifications. In this context, we utilized WGA sequencing data as a negative control, enabling stringent filtering of high-confidence methylation sites, thereby effectively eliminating a significant number of potential false-positive sites. The high conservation and motif coverage rates of these methylated sites further validated the effectiveness of our methodology. Despite the filtering, however, persistent noise remained in the detection of 4mC modifications due to interference from adjacent 6mA signals. Additionally, a small fraction of methylation sites identified did not match known motifs and were only marginally above filtering thresholds (Supplementary Fig. S7). To ensure high analytical accuracy, our subsequent analysis focused exclusively on high-quality methylated sites associated with well-characterized motifs for 6mA, 5mC, and 4mC modifications. The authenticity of these motifs was further reinforced through cross-validation with WGA sequencing data and confirmation of the corresponding DNA methyltransferase genes. Taken together, we minimized the false positives in the derived methylation profiles to the greatest extent by implementing the rigorous filtering criteria and multiple evaluations, supported by WGA sequencing data.

### 4.3 Demethylation of CAYNNNNNRTG motif as a robust epigenetic response to starvation, potentially affecting gene expression and metabolic processes

One interesting question in bacterial epigenetics is how DNA methylation patterns respond to prolonged starvation stress. Previous studies have characterized stress-responsive methylation patterns in various bacterial species. For example, survival of *Escherichia coli* under antibiotic stress has been linked to 6mA methylation of the GATC motif (Cohen et al., 2016). Similarly, in *Salmonella enterica*, Zhang et al. (2023) demonstrated that methylation of the GATC motif contributes to oxidative stress adaptation through genome-wide knockout of methyltransferase genes. In *Lacticaseibacillus paracasei*, a combination of gene knockout and multi-omics approaches confirmed that 6mA methylation of the ACRCAG motif plays a regulatory role in adaptation to ethanol and osmotic stress (Yan et al., 2023). These studies collectively suggest that specific 6mA-associated methylation motifs are critical for bacterial stress adaptation. In line with these findings, our study identified the 6mA-associated CAYNNNNNRTG motif as a key epigenetic element in response to prolonged starvation. Notably, this motif was the only one to exhibit a consistent decline in methylation levels under starvation, regardless of temperature. Moreover, the methylation sites that were significantly downregulated under starvation at both temperatures were predominantly associated with this motif. This starvation-induced demethylation pattern resembles previously observed methylation reductions under low-temperature stress (Bu et al., 2025). Functional enrichment analysis revealed that these CAYNNNNNRTG-associated demethylated sites were mainly located within genes involved in tRNA aminoacylation and molecular metabolism.

DNA methylation-dependent gene regulation has already been implicated in bacterial stress adaptation (Ghosh et al., 2020; Motta et al., 2015). While this regulatory mechanism is often associated with methylation in promoter regions, recent evidence highlights a regulatory role for methylation within coding sequence (CDS) as well (Kahramanoglou et al., 2012; Kumar et al., 2018). Based on our results, we propose that the CAYNNNNNRTG-specific demethylation may represent a coordinated regulatory mechanism modulating metabolic and translational pathways during bacterial adaptation to starvation. This is consistent with known physiological responses of bacteria under stress, where cells enter the dormant-like state characterized by global reductions in gene expression and metabolic activity, retaining only essential cellular functions (Harms et al., 2016; Sachidanandham and Yew-Hoong Gin, 2009; Zhang et al., 2014). For example, a sharp decrease in translation rate has been reported in mycobacteria under starvation conditions (Li et al., 2022). In starved *E. coli*, levels of translation-associated proteins were also significantly reduced (Shi et al., 2021). Although metabolically active, starved bacteria function at substantially reduced metabolic levels, relying on limited, specific pathways for survival (Greening et al., 2019; Lempp et al., 2020). This includes the upregulation of stress-related oxidoreductases, as seen in dormant *E. coli* populations amid global metabolic downshifts (Clegg et al., 2006). Furthermore, mathematical modeling studies support the notion that nutrient deprivation leads to a slowdown or cessation of metabolism (Nev and van den Berg, 2018).

### 4.4 GCAGA motif exhibits a temperature-dependent starvation response, likely regulating DNA replication

In addition to the starvation-induced demethylation of the CAYNNNNNRTG motif, we also observed a distinct, motif-specific methylation response associated with temperature stress under prolonged starvation. Our experimental design, which examined methylation patterns under starvation at two temperatures associated with different growth dynamics, allowed us to dissect temperature-dependent methylation changes. Notably, we observed significant methylation changes primarily associated with another 6mC-modified GCAGA motif, which was enriched within genes involved in DNA replication. This finding suggests a potential role for GCAGA in mediating temperature-specific adaptive responses under nutrient-deprived conditions. Importantly, these results underscore the motif-specific nature of bacterial methylation responses, with distinct motifs acting as epigenetic switches that respond to different environmental stressors.

The role of temperature in bacterial starvation adaption has been already well described in prior studies (Arana et al., 2010; Auty et al., 2022). Dormancy is a widely recognized survival strategy under starvation stress for bacteria including *F. columnare*, while alternative adaptive strategies such as reduced growth rates have also been proposed (Biselli et al., 2020; Gray et al., 2019). Our observation of temperature-specific methylation patterns targeting replication-associated genes suggests an additional layer of complexity, where temperature-induced epigenetic regulation via the GCAGA motif may contribute to controlling gene expression involved in bacterial replication processes. Taken together, our results provide evidence for the existence of motif-specific methylation regulation, possibly allowing bacteria to fine-tune gene expression and physiological responses under different combinations of starvation and temperature stresses.

### 4.5 Potential link between epigenetic changes and starvation-induced colony morphology and virulence

It is also noteworthy that after a 10-month period of starvation at different temperatures, *F. columnare* exhibited a consistent alteration in colony morphology from the original rhizoid form to a smooth phenotype. The rhizoid morphology was associated with gliding motility, a mechanical movement process independent of flagella and pili (Chang et al., 1984; Penttinen et al., 2018). In *Flavobacterium* spp., this gliding motility has been well characterized and involves proteins in the Type IX secretion system (T9SS), which transport motility-associated adhesins to the cell surface (McBride and Nakane, 2015; Penttinen et al., 2018). We hypothesize that under prolonged starvation, reduced translational activity may impair the production of these gliding-related proteins in *F. columnare*, contributing to the observed morphological transition. Importantly, the T9SS in *F. columnare* is also implicated in the secretion of virulence factors, thus closely linked to bacterial virulence (Dumpala et al., 2010). Additionally, we observed significantly reduced methylation levels in stress-related pathway related to siderophore transmembrane transport, under starvation at both tested temperatures. This process not only supplies bacterial cells with iron essential for growth and metabolism but is also associated with bacterial virulence (Conrad et al., 2022). Therefore, we propose that starvation-induced DNA demethylation may downregulate virulence-related pathways in *F. columnare*. This is consistent with previous findings that the smooth morphology exhibits lower virulence compared to the rhizoid morphology (Kunttu et al., 2009; Laanto et al., 2014).

## 5. Conclusion

In conclusion, Nanopore R10.4.1 sequencing offers an exceptional opportunity to investigate DNA methylation at single-base resolution in aquatic bacteria, albeit with certain limitations in the detection of 4mC modifications. Our methylation-based analysis demonstrated that while the genomic sequence of *F. columnare* remained remarkably stable, its DNA methylation landscape exhibited motif-specific plasticity under prolonged starvation. The specific demethylation pattern of the CAYNNNNNRTG motif represented a robust epigenetic signature in response to starvation stress, particularly evident in genes related to translation and molecular metabolism. Temperature also induced distinct starvation-related methylation changes, notably involving the GCAGA motif, where DNA replication gene-related sites displayed significant methylation differences between room temperature (22 °C) and optimal growth temperature (28 **°**C). These findings provide evidence supporting epigenetic regulation as one of the fundamental mechanisms enabling bacterial adaptation to prolonged nutrient limitation. Furthermore, this study serves as a valuable reference for future microbiological research employing ONT for bacterial methylation analysis.

## Declaration of Competing Interest

The authors declare that they have no known competing financial interests or personal relationships that could have appeared to influence the work reported in this paper.

## Funding

This study was supported by the APRC-CityU New Research Initiatives/Infrastructure Support from Central (9610574, 7006064), and Early career scheme (project number 9048294) from the Hong Kong Research Grant Council.

## Author contributions

**Zhang Yuxuan:** Conceptualization, Formal analysis, Investigation, Methodology, Visualization, Writing–original draft; Writing–review and editing. **Shao Yanwen:** Conceptualization, Formal analysis, Investigation, Writing–review and editing. **Gao Shengnan:** Data curation, Writing–review and editing. **Li Runsheng:** Conceptualization, Methodology, Supervision, Writing–review and editing. **Cai Wenlong:** Conceptualization, Funding acquisition, Methodology, Supervision, Writing–review and editing.

## Data availability

All the sequencing data and genome assemblies have been deposited in NCBI BioProject with the accession number PRJNA1247036.

## Supplementary materials

**Figure S1:** Colony morphology change caused by long-term starvation.

**Figure S2:** 1% Agarose gel electrophoresis of genomic DNA extracted from Fc-con, Fc-22, and Fc-28.

**Figure S3:** Filtering of raw base modifications in the native DNA of Fc-22 and Fc-28 output by Dorado.

**Figure S4:** Box plots showing the comparison of raw 6mA, 5mC, and 4C methylation levels between WGS and WGA for Fc-con, Fc-22, and Fc-28.

**Figure S5:** Functional analysis of genes containing each methylated motif.

**Figure S6:** Comparison of methylation levels and difference indices for each motif between WGS and WGA sequencing.

**Figure S7:** Motif coverage among the identified high-confidence methylation sites.

**Figure S8:** Differential reduction patterns in methylation levels in Fc-28 and Fc-22, compared to Fc-con.

**Table S1:** Statistics of sequenceing data for Flavobacterium columnare.

**Table S2:** Statistics of SNPs and INDELs identified in Flavobacterium columnare under starvation conditions.

**Table S3:** Number of sites with raw modification signals detected in native DNA of Fc-con, Fc-22, and Fc-28.

**Table S4:** False positive rate (FDR) and number of remaining sites under different cutoffs for 6mA methylation levels.

**Table S5:** False positive rate (FDR) and number of remaining sites under different cutoffs for 5mC methylation levels.

**Table S6:** False positive rate (FDR) and number of remaining sites under different cutoffs for 4mC methylation levels.

**Table S7:** Motifs identified based on high-confidence methylated sites using STREME.

**Table S8:** Motifs identified based on high-confidence methylated sites using Modkit

**Table S9:** Abundance of methylated motifs in Fc-con, Fc-22, and Fc-28. The methylated sites are marked with underscores.

**Table S10:** MTases assignment to identified methylated motifs by BLASTP against REBASE and Nanomotif.

**Table S11:** Number of motif-associated methylated sites distributed in different genomic features.

## Notes

### Competing Interest Statement

The authors have declared no competing interest.

