## Supplemental Figures for "Prolonged Starvation Drives Epigenetic Remodeling: Insights from DNA Methylation Profiling in the Aquatic Pathogen *Flavobacterium columnare*"

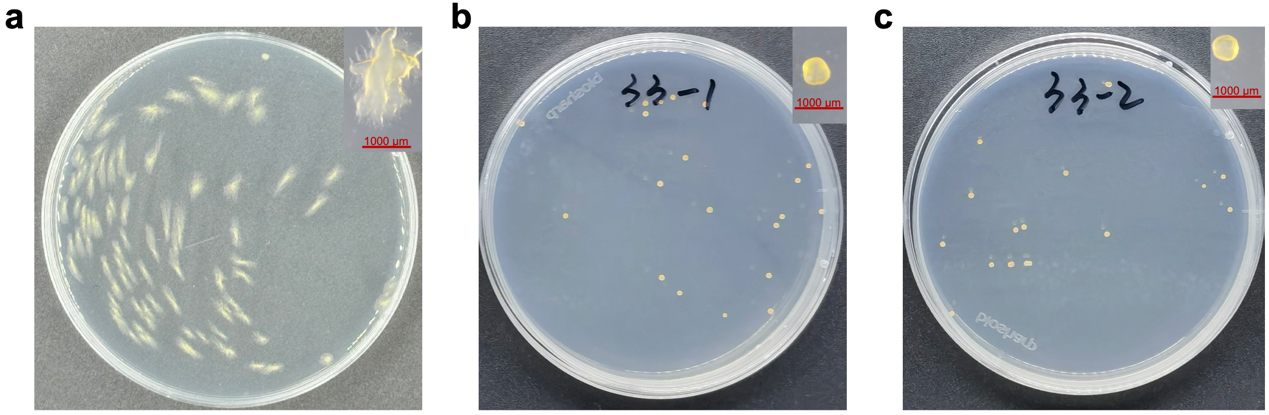

**Figure S1.** Colony morphology change caused by long-term starvation. (**a**) Colony morphology of Fc-con under the nutrient-rich condition. (**b**) Colony morphology of Fc-28 under 28 **°C** nutrient-deficient condition. (**c**) Colony morphology of Fc-22 under 22 **°C nutrient-deficient condition. The top-right figures show colony morphologies under stereomicroscopy at 10** × magnification.

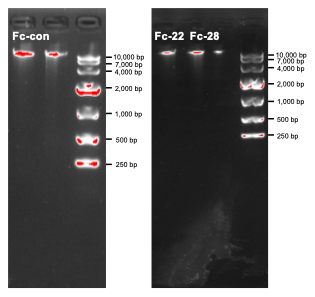

**Figure S2.** 1% Agarose gel electrophoresis of genomic DNA extracted from Fc-con, Fc-22, and Fc-28.

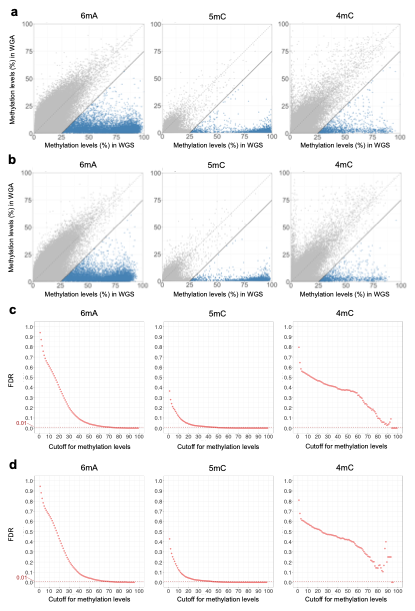

**Figure S3.** Filtering of raw base modifications in the native DNA of Fc-22 and Fc-28 output by Dorado. (**a-b**) Filtering based on the difference in methylation levels between WGA and WGS sequencing for Fc-22 (a) and Fc-28 (b). The points represent individual base modifications. Most base modifications are distributed around the diagonal, indicating highly similar methylation levels between WGS and WGA. The blue points represent high-confidence base modifications filtered using the criterion "WGS methylation level - WGA methylation level > 25%". (**c-d**) False discovery rates (FDR) under filtering based solely on methylation levels in WGS sequencing for Fc-22 (c) and Fc-28 (d). Methylation levels required for FDR < 0.01 are significantly high.

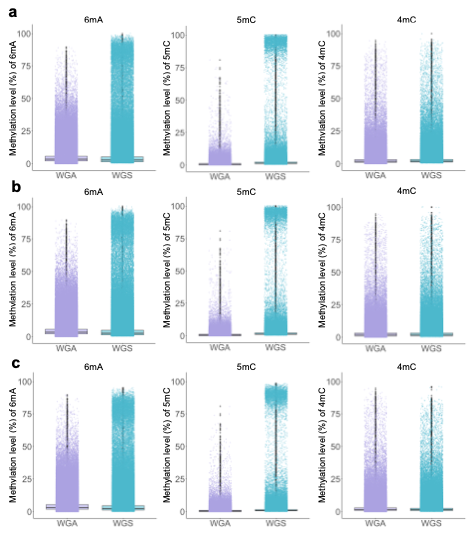

**Figure S4.** Box plots showing the comparison of raw 6mA, 5mC, and 4C methylation levels between WGS and WGA for Fc-con (**a**), Fc-22 (**b**), and Fc-28 (**c**).

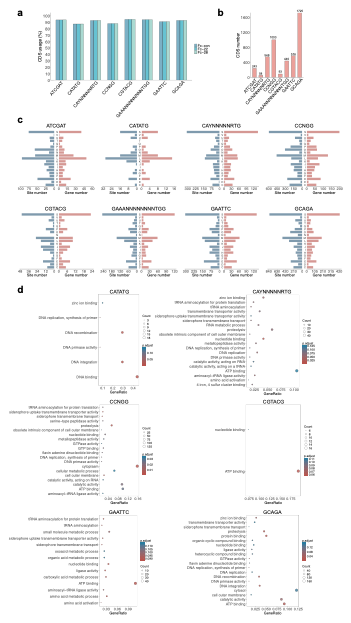

**Figure S5.** Functional analysis of genes containing each methylated motif. (**a**) CDS utilization rates of each methylated motif in Fc-con, Fc-22, and Fc-28. (**b**) The number of genes containing each methylated motif. Only the sequence from one strand is displayed for non-palindromic bipartite motifs. (**c**) COG classification of genes containing each motif. COG functional categories: [C], Energy production and conversion; [D], Cell cycle control, cell division, chromosome partitioning; [E], Amino acid transport and metabolism; [F], Nucleotide transport and metabolism; [G], Carbohydrate transport and metabolism; [H], Coenzyme transport and metabolism; [I], Lipid transport and metabolism; [J], Translation, ribosomal structure and biogenesis; [K], Transcription; [L], Replication, recombination and repair; [M], Cell wall/membrane/envelope biogenesis; [O], Posttranslational modification, protein turnover, chaperones; [P], Inorganic ion transport and metabolism; [S], Function unknown; [T], Signal transduction mechanisms; [U], Intracellular secretion, trafficking, and vesicular transport; [V], Defense mechanisms. (**d**) GO enrichment of genes containing each motif. When the number of significant GO terms (Benjamini-Hochberg (BH)-adjusted *p*-value < 0.05) exceeds 20, only the top 20 GO terms are displayed. No significantly enriched GO terms were identified for genes containing the ATCGAT and GAAANNNNNNNNTGG motifs

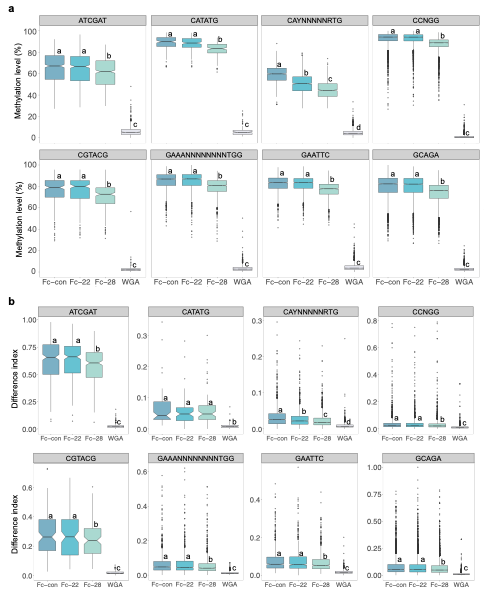

**Figure S6.** Comparison of methylation levels and difference indices for each motif between WGS and WGA sequencing. (**a**) Comparison of methylation levels obtained using Dorado. (**b**) Comparison of difference indices obtained using Hammerhead. Different letters above the boxes indicate significant differences (Wilcoxon signed-rank test *p* < 0.01).

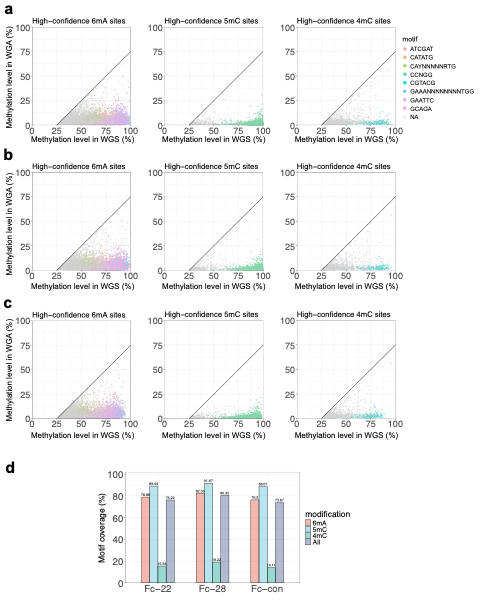

**Figure S7.** Motif coverage among the identified high-confidence methylation sites. (**a-c**) Distribution of motif-associated sites among high-confidence 6mA, 5mC, and 4mC modified sites in Fc-con (a), Fc-22(b), and Fc-28 (c). (**d**) Statistics on motif coverage in high-confidence 6mA, 5mC, and 4mC modified sites.

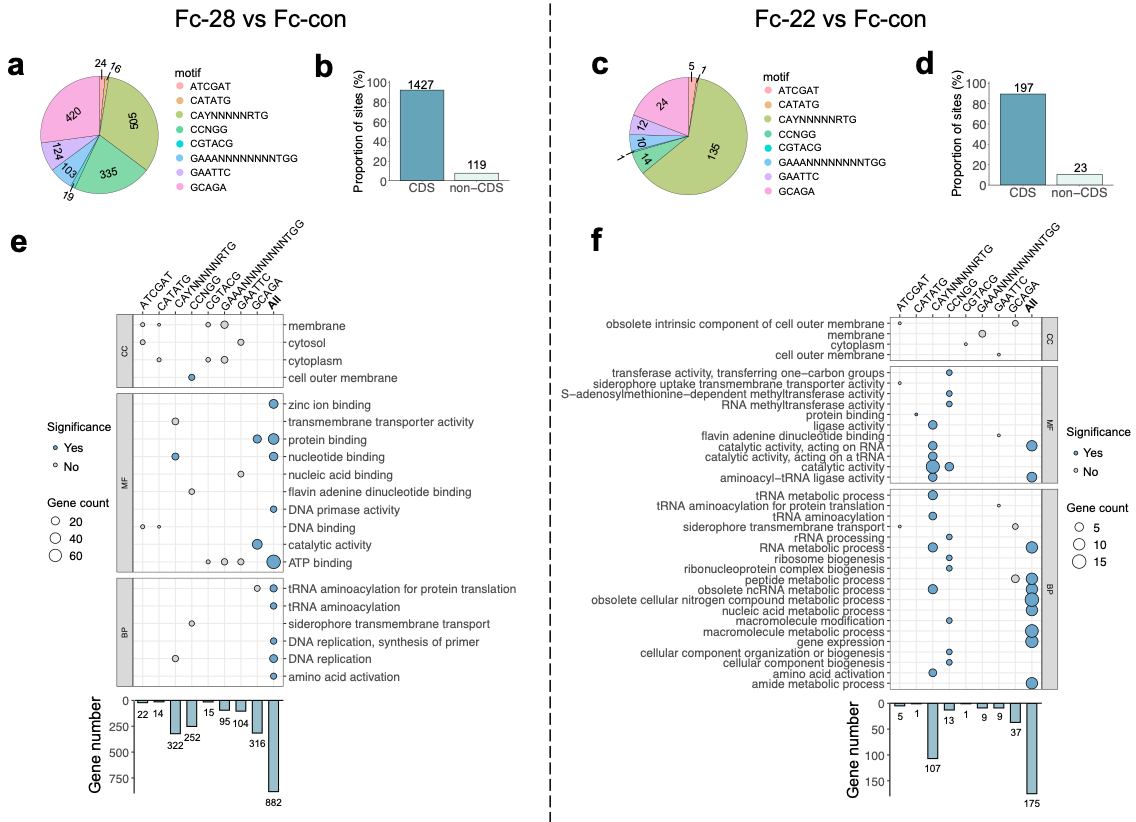

**Figure S8.** Differential reduction patterns in methylation levels in Fc-28 and Fc-22, compared to Fc-con. (**a-b**) Proportion of motifs (a) and CDS usage (b) associated with significantly downregulated sites in Fc-28, compared to Fc-con. The numbers in the pie chart and on the bars indicate the corresponding site counts. (**c-d**) Proportion of motifs (c) and CDS usage (d) associated with significantly downregulated sites in Fc-22, compared to Fc-con. (**e-f**) Bar plots display the number of CDSs containing significantly downregulated sites for each motif in Fc-28 (e) and Fc-22 (f). Dot plots show the GO enrichment analysis of genes containing significantly downregulated sites for each motif in Fc-28 (e) and Fc-22 (f). When more than 3 significant GO terms (Benjamini-Hochberg (BH)-adjusted *p*-value < 0.05) were identified, the top 10 are shown, and when fewer than 3 significant GO terms were identified, the top 3 are displayed. Blue and gray dots represent significant and non-significant enrichment, respectively.
